# Individual based genetic analyses support asexual hydrochory dispersal in *Zostera noltei*

**DOI:** 10.1101/340554

**Authors:** Buga Berković, Nelson Coelho, Licínia Gouveia, Ester A. Serrão, Filipe Alberto

**Affiliations:** Centro de Ciências do Mar, Universidade do Algarve, Faro, Portugal; Department of Biological Sciences, University of Wisconsin – Milwaukee, Milwaukee, Wisconsin, United States of America; Department of Computational and Systems Biology, School of Medicine, University of Pittsburgh, Pennsylvania, United States of America

**Keywords:** asexual dispersal, hydrochory, clonal reproduction, seagrass, microsatellites, *Zostera noltei*

## Abstract

Dispersal beyond the local patch in clonal plants was typically thought to result from sexual reproduction via seed dispersal. However, evidence for the separation, transport by water, and re-establishment of asexual propagules (asexual hydrochory) is mounting suggesting other important means of dispersal in aquatic plants. Using an unprecedented sampling size and microsatellite genetic identification, we describe the distribution of seagrass clones along tens of km within a coastal lagoon in Southern Portugal. Our spatially explicit individual-based sampling design covered 84 km^2^ and collected 3 185 *Zostera noltei* ramets from 803 sites. We estimated clone age, assuming rhizome elongation as the only mechanism of clone spread, and contrasted it with paleo-oceanographic sea level change. We also studied the association between a source of disturbance and the location of large clones. A total of 16 clones were sampled more than 10 times and the most abundant one was sampled 59 times. The largest distance between two samples from the same clone was 26.4 km and a total of 58 and 10 clones were sampled across more than 2 and 10 km, respectively. The number of extremely large clone sizes, and their old ages when assuming the rhizome elongation as the single causal mechanism, suggests other processes are behind the span of these clones. We discuss how the dispersal of vegetative fragments in a stepping-stone manner might have produced this pattern. We found higher probabilities to sample large clones away from the lagoon inlet, considered a source of disturbance. This study corroborates previous experiments on the success of transport and re-establishment of asexual fragments and supports the hypothesis that asexual hydrochory is responsible for the extent of these clones.

## Introduction

Clonal propagation is widespread throughout the Tree of Life, including multicellular eukaryotes, in 45% of vascular plant families [1] and over 70% of animal phyla [2]. Most eukaryotic clonal organisms are also capable of reproducing sexually, and will carry out one mode of reproduction or the other according to the circumstances [3–5]. Dispersal is one of many important life history traits associated with sexual reproduction: sexually-derived bodies (such as gametes, pollen, zygotes, seeds, larvae) are often thought of as the relevant dispersal propagules mediating gene flow within and between populations. On the other hand, asexual reproduction was for long thought to play a role constrained to the population spatial limits, such as resource foraging through clonal extension, physical and physiological integration and population maintenance under mate limitation [6]. This clonal extension, or growth, is equivalent to asexual dispersal because it constitutes movement of genes away from zygote natal site [7]. In clonal plants this is achieved by creeping rhizomes, stolons, or roots. We note that this type of asexual dispersal occurs with the organism never leaving the substrate, although the physical connection between modules might be kept or lost. Thus, the rate of asexual dispersal through this mechanism is tightly associated with specific clonal growth rates which are generally slow.

However, in aquatic organisms, a much faster asexual dispersal mechanism is the aquatic transport of asexual propagules, a form of asexual hydrochory (AH). Evidence for AH is mounting calling attention to the role of asexual dispersal at spatial scales beyond the local population [8,9]. AH occurs when an asexual propagule is fragmented and temporally escapes its sessile dependency on the substrate. This type of propagule dispersal is a three-stage process: a *separation* from the established clone and from the substrate, followed by a *transport* period, and finally, the *establishment* in a new location. The potential to produce viable vegetative fragments has been observed and demonstrated in a number of animal and plant species [10–15]. The water transport phase of asexual dispersal in nature is often difficult to track, due to practical limitations to observe or experimentally test propagule dispersal, particularly beyond the local patch. Settlement of the dispersed asexual propagules has been confirmed in several taxa: plants in lentic habitats [16], in streams [12,17,18], marine macrophytes [19,20], freshwater bryozoans [21], moss [22], sea anemones [23], corals [14,24] and sponges [11], among others. Post dispersal establishment of fragments of submerged freshwater macrophytes is well supported [12,25,26]. Studies which looked into this process in marine macrophytes reported successful establishment [19,20,27–30].

Seagrasses are marine plants relying on both clonal growth and sexual reproduction to extend their coverage in the habitat or to occupy new ones. Dispersal distance by sexual propagules is often limited when they are on their own in the marine environment [31], although dispersal distances can be orders of magnitude longer when seeds are transported by reproductive structures or fragments floating in the current [6,31–34]. Perhaps because of this, studies on seagrass dispersal disproportionately focus their attention on seed dispersal, while asexual dispersal is mainly associated with rhizome extension. However, evidence for the different elements necessary for AH in seagrasses is mounting. Production of seagrass fragments is common across species regardless of their size or habitat and can be a natural or human-induced process [35–37]. The transport of such fragments was reported for the first time in 1908 [38] and has since been suggested many times [39–44] or as a transport vector for seeds [34]. Using genetic markers to characterize clonality, Arnaud-Haond et al. [45] suggested that fragment dispersal could explain the distribution of very large clones, along areas where seagrasses could not have persisted during the low sea levels of the most recent glacial period. While recently, the largest extension covered by a single seagrass clone was found in the Indian River Lagoon, Florida, for *Thalassia testudinum* [9]. The mechanism proposed for the wide extension of this clone was AH. Across these studies, evidence is still lacking for the success of all three steps necessary for AH in the same species. Berković et al. [46] study demonstrated long viability of seagrass shoots, and seeds carried in them, during the transport phase of *Z. noltei* fragments. In a follow up study, Berković [47] showed biologically relevant post-dispersal successful settlement in *Z. noltei*, ranging between 30 and 100% depending on the size of the fragment, the time spent floating, and the time after settlement. These studies provided the motivation to find a spatial signature of AH that relies on genetic identification of clones.

The use of hyper-variable genetic markers (e.g. microsatellites) in the last 15 years has increased our knowledge on the spatial patterns of clonal structure [48]. Unbiased genetic identification of the same clone sampled across distant locations has been explained as a result of rhizome elongation [45,49,50]. Clone age estimation based on species’ rhizome elongation rates might lead to overestimated ages if AH is overlooked. Using this age estimation method, some seagrass species are mentioned among the oldest organisms on the planet [45,49]. However, when estimating clone age using this approach, rarely has the hypothesis of AH been adequately considered using experimental studies, possibly avoiding age overestimation by orders of magnitude.

Most sampling designs used in population genetics are compromised in their capacity to reveal AH. Many studies have a population-based spatial scale with “populations” arbitrarily defined and sampled in clusters, leaving a large proportion of unsampled habitat in between. On the other hand, when the focus is on within-population processes, as in fine spatial scale genetic structure analysis, sampling densities are higher but the spatial extent of the sample is limited. To circumvent these limitations one should use an individual-based, spatially-explicit sampling design, which is common in landscape genetics [51]. In this design, sample units are collected randomly or stratified over the entire habitat under study. Such strategy, if including large enough sample size, increases the chance of sampling multiple ramets of the same clone spread over large distances in a continuous or disjunctive mode. This sampling design has only been applied with limited spatial extent in studies of sessile clonal organisms.

In this study, we used an individual-based sampling design, across an unprecedented spatial extent, to study the clonal structure of the seagrass *Zostera noltei* using microsatellite markers. Given previous experimental work supporting the capacity for AH in this species [46,47], we predicted finding large spatial extent of clones within the Ria Formosa.

## Materials and methods

### Study site

This study was carried out in the Ria Formosa lagoon, Portugal (37°N 8°W). This intertidal lagoon extends roughly 55 km along the mainland and is 6 km across its widest point, with an average depth of about 2 m and low fresh water input. Separated from the ocean by five islands and two peninsulas, it consists of a complex set of channels, mudflats, saltmarshes and highly dynamic sand barrier islands. Intertidal mudflats are inhabited by the seagrass *Z. noltei*, whereas subtidal areas are habitat to two other seagrass species, *Cymodocea nodosa* and *Zostera marina*.

### Study species

*Zostera noltei* is a small seagrass species distributed along central/southern Europe and NW Africa [52]. In the Mediterranean it is mostly found in the shallow subtidal zone, with lower shoot density, while being intertidal in most Atlantic meadows with high density [53–55]. *Z. noltei* is the dominant seagrass species in the Ria Formosa lagoon, covering over 13 km^2^ of the intertidal area [56]. The species shows highly variable growth rates, but it is generally considered a fast growing species. Alexandre et al. [57] reported frequent flowering and high production of seeds in the Ria Formosa for this monoecious species. Seeds can be released directly from the plant or can be transported attached to the flowering shoot [58]. Seed transport distance ranges from tens of cm, when released from an attached parental shoot, to a couple thousand kilometers when transported by buoyant detached shoots, assuming unidirectional current flow [46]. Detached positively buoyant fragments keep producing new shoots and can carry maturing seeds for more than 50 days [46]. An experimental study on post-dispersal settlement success, after a floating period of up to four weeks, demonstrated that these fragments once entangled are quickly buried and keep growing [47].

### Sample collection

We randomly selected 1 000 sampling coordinates within a 13 km^2^ area of *Z. noltei* meadows in the Ria Formosa, based on an available georeferenced distribution across more than 84 km^2^ of the lagoon [56]. Due to logistic constrains only 899 of these plots were visited by boat, kayak and walking within the intricate saltmarsh channels. *Z. noltei* was found at 803 of these coordinates. At each plot, 4 sample units were collected at the vertices of a 4 m^2^ quadrat. Each individual sample unit consisted of a single ramet with three to four shoots connected by a horizontal rhizome. Thus, a total of 3 212 individual ramet sample units were sampled. Back in the laboratory, samples were carefully washed in fresh water, dried on paper and stored dry with silica gel.

### Genetic analysis of clone identity

Genomic DNA for all ramets sampled was isolated from silica dried tissue (5-10 mg) using the CTAB method [59]. All *Z. noltei* samples were genotyped for nine microsatellite loci following Coyer et al. [60]. We used fluorochrome-labelled primers on a GeneAmp 2700 thermocycler (Applied Biosystems, Foster City, CA, USA) and an ABI PRISM 3130xl DNA analyser (Applied Biosystems) was used to determine the size of amplification products (i.e., microsatellite alleles). We scored raw fragment sizes using STRand v.2.4.59 (http://www.vgl.ucdavis.edu/informatics/strand.php), and binned into allele classes using the R (R core team, 2016) package MsatAllele [61]. The probability that sampled ramets with observed identical multilocus genotypes (MLGs) were different genets originated by distinct sexual reproduction events, *psex* [62,63], was estimated using custom R code.

### Estimating clonal age using rhizome elongation rates

Typically, a seagrass individual’s age is estimated assuming that the sampled spatial extent of a clone is the result of rhizome elongation alone [45,49]. We used a similar strategy, but iteratively corrected age estimates by combining sampling locations with spatial explicit sea-level data for the time the clone would have been initiated. Once clonal assignment was determined, we determined the longest “as-the-crow-flies” distance between clonemates (ramets belonging to the same genet). This distance between a pair of clonemates A and B is likely an underestimate (i.e., sampled) of clone’s span. We then conservatively assumed that the clone initiated growth at the middle point (O_0_) in a line between A and B. An initial estimate of clone age (t_0_) was calculated by dividing the distance from O_0_ to A (half the clone’s span) by *Z. noltei* growth rate. We used a mean rate of rhizome elongation of 68 cm/year from Marbà and Duarte [64] review, including different areas and seasons to reflect changes in habitat and climate through time. The estimated age of these clones was then matched with sea-level change reconstruction using paleoceanographic data over the past 20 000 years for Southern Portugal [65] and current bathymetry. Clone age estimated above allowed us to determine if at the inferred time of clone initiation O_0_ was submerged or on land. If the latter was true, we reiterated the clone aging process by moving O_0_ to the closet submerged location at the date corresponding with clone age. The iteration continued until we found a submerged O_0_ or we runned out of paleoceanographic data (details in S1 File and S1 Fig).

To further characterize the spatial distribution of *Z. noltei* clones within the Ria Formosa, we estimated for each plot the average probability of sampling an MLG that was present in the sample more than 5, 10 and 15 times (*Pc5, Pc10* and *Pc15*, respectively). Note that *Pc5* represents the chance of sampling a clone that was found at least in two plots, because only four ramets were sampled per plot (average shortest distance between plots: 96 m). Finally, we investigated how *Pc* changed with distance from the lagoon’s barrier island system, a putative source of disturbance [66] (details in S2 File).

## Results

The genetic analysis of 3 185 *Zostera noltei* ramets revealed 1 999 unique MLGs corresponding to a relatively high genotypic richness (MLGs/N=R=0.63), with mean allelic richness per locus A=15.0, observed heterozygosity Ho=0.62 and probability of sexual re-sampling, *Psex*, always <0.001. Out of the unique MLGs, 504 were sampled more than once. A total of 16 MLGs were sampled more than 10 times and the most abundant MLG (ID 1886) was sampled 59 times (Table 1, Fig 1 c). The largest distance between two samples with identical MLG was 26.4 km (Fig 1 b). A total of 58 MLGs were sampled across more than 2 km and out of those 10 MLGs were sampled across more than 10 km. An example of the large spatial extent of four these clones is shown in Fig 2 a. To illustrate the necessary power to detect asexual hydrochory, we calculated the proportion of ramet pairwise combinations that shared the same MLG and plotted it as a function of the distance between sampled ramets (Fig 2). Although many clones were extended across several kilometers, the proportion of pairwise samples that captured clonemates decreased with distance and was generally small, 1.62% and 0.12%, at 50 m and 30 km distance classes, respectively (Fig 2).

**Fig 1.**
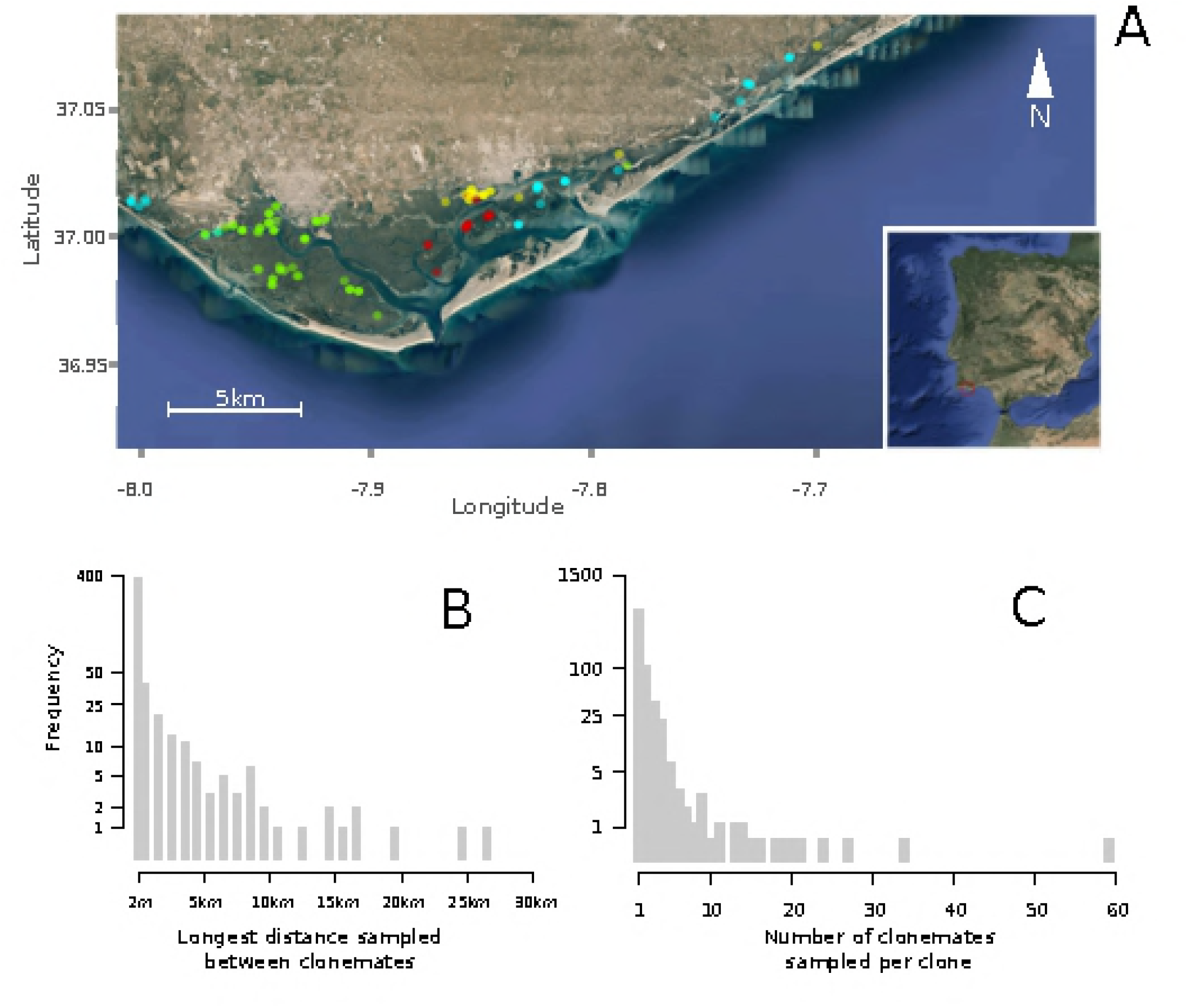
Clonal spatial structure of *Zostera noltei* in Ria Formosa, Portugal. Panel A shows the distribution of four large *Zostera noltei* clones. The circles represent sampling plots and the colors used show where the same clone was sampled. The most opaque colors indicate that all four ramets sampled in a plot had that particular clone. Histograms show the distributions of the longest distance sampled between clonemates, panel B, and the number of clonemates sampled per clone, panels c.

**Fig 2.**
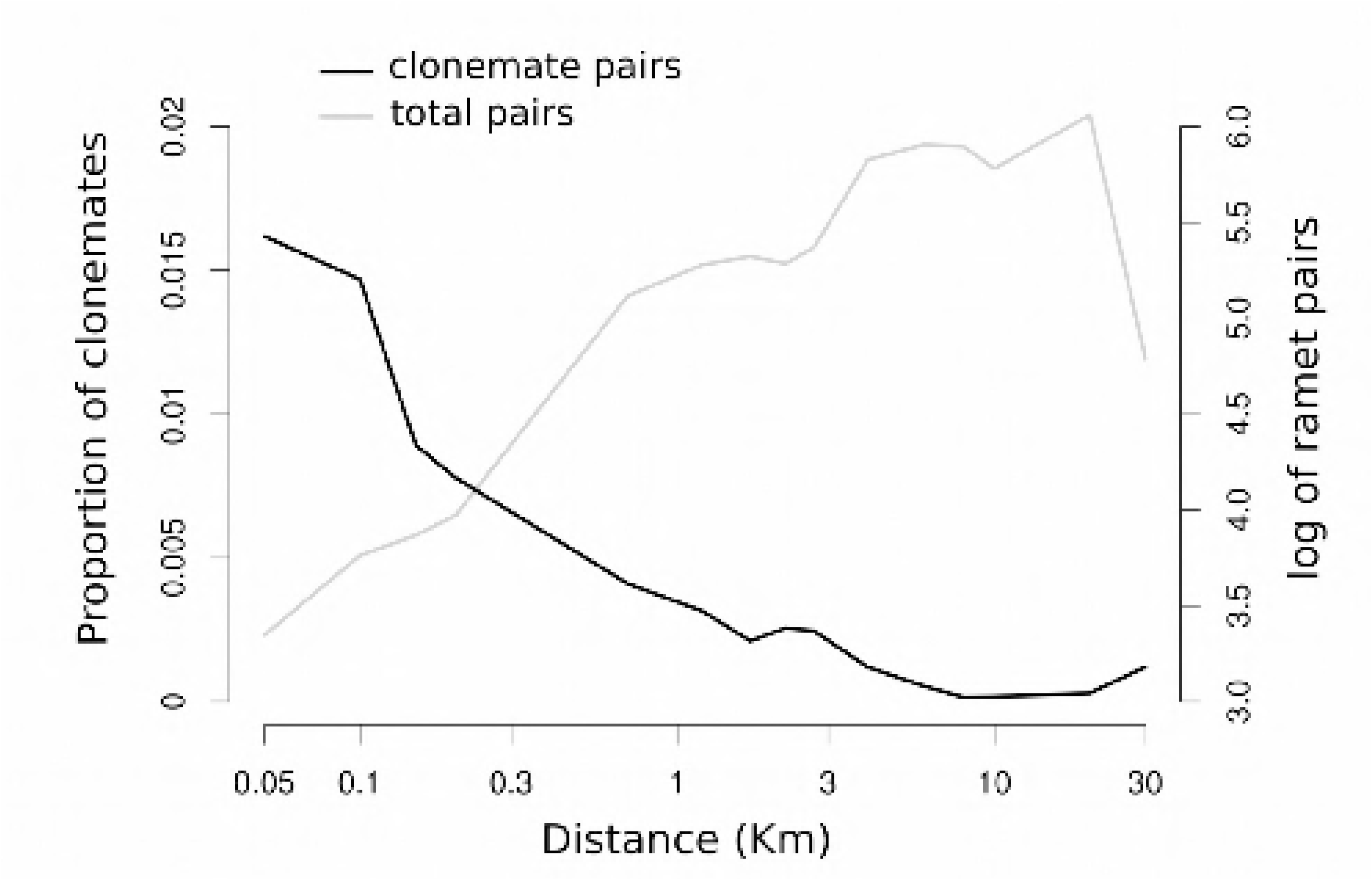
Change with distance between sample units in the proportion of clonemates pairs. This figure illustrates the need for large individual based sample designs to reveal large clonal spans. The black line indicate the proportion of ramet pairs, sampled at different spatial distance classes, that showed the same *Z. noltei* genotype (clonemates). The gray line shows the large total number of ramet pairwise combinations that were compared at each distance class (log10 scale is being used).

**Table 1.** Age estimation for the ten largest clones of *Zostera noltei* assuming no asexual hydrochory. N – number of clonemates sampled; distance – maximum distance sampled between two clonemates; O_1_ through O_4_ – shortest distance to the estimated point of origin; t_0_ through t_3_ – estimated age of the clone assuming rhizome elongation from the point of origin at different sea levels (APPENDIX A); * – final age estimation after correcting for sea level; n.a. – unavailable sea level data after this point, clone older than last age estimated. Data is also shown for one clone of the seagrass *Cymodocea nodosa* in the Ria Formosa lagoon (reanalyzed from Alberto et al. 2008)

| Species | Clone | N | Distance (km) | O<sub>0</sub> (km) | t<sub>0</sub> (y) | O<sub>1</sub>(km) | t<sub>1</sub> (y) | O<sub>2</sub>(km) | t<sub>2</sub> (y) | O<sub>3</sub>(km) | t<sub>3</sub> (y) | O<sub>4</sub>(km) |
| --- | --- | --- | --- | --- | --- | --- | --- | --- | --- | --- | --- | --- |
| *Zostera noltei* | 1 259 | 34 | 26.4 | 13.2 | 19 421 | 22.5 | 33 088 | n.a. |  |  |  |  |
|  | 120 | 6 | 24.1 | 12.0 | 17 710 | 22 | 32 352 | n.a. |  |  |  |  |
|  | 896 | 11 | 19.6 | 9.8 | 14 389 | 19 | 27 941 | n.a. |  |  |  |  |
|  | 1 886 | 59 | 16.7 | 8.4 | 12 306 | 13 | 19 117 | 16.5 | 24 264 | n.a. |  |  |
|  | 1 480 | 9 | 16.2 | 8.1 | 11 879 | 15 | 22 058 | n.a. |  |  |  |  |
|  | 737 | 19 | 15.9 | 8 | 11 710 | 15 | 22 058 | n.a. |  |  |  |  |
|  | 1 386 | 2 | 15.0 | 7.5 | 11 001 | 10 | 14 705 | 12 | 17 647 | 13 | 19 117* |  |
|  | 44 | 3 | 14.1 | 7 | 10 368 | 11 | 16 176 | 12.5 | 18 382 | 13 | 19 117* |  |
|  | 1 524 | 13 | 12.3 | 6.2 | 9 070 | 10 | 14 705 | 13 | 19 117 | 15 | 22 058 | n.a. |
|  | 707 | 6 | 10.8 | 5.4 | 7 993 | 10 | 14 705 | 12.5 | 18 382 | 13.5 | 19 852* |  |
| *Cymodocea nodosa* | 1 | 176 | 42.9 | 21.4 | 53635 |  |  |  |  |  |  |  |

Using the rhizome elongation rate method, corrected for sea-level change, we estimated that among the 10 largest clones three could be approximately 20 000 years old, while the remaining seven to be older than 20 000 years. Further iterations of our age estimation process beyond 20 000 years before present (YBP) were not possible, due to the temporal extent of available paleoceanographic data for this region. However, the age of these older clones would have to be at least equal or older than the range found at the point where we could no longer proceed (22 058 to 33 088 years, Table 1).

The analysis of spatial distribution of large *Z. noltei* clones within the lagoon revealed higher probability to sample large clones (*Pc*) further away from the lagoon barrier islands (Fig.3 a). The average clonal probability *Pc5* per plot was 0.05 at 400 m away from the barrier island and increased to 0.35 at 4 300 m, with a linear increase of 0.075 per km. This pattern did not change with the number of clonemates used to estimate *Pc* (5, 10 or 15, Fig 3 b).

**Fig 3.**
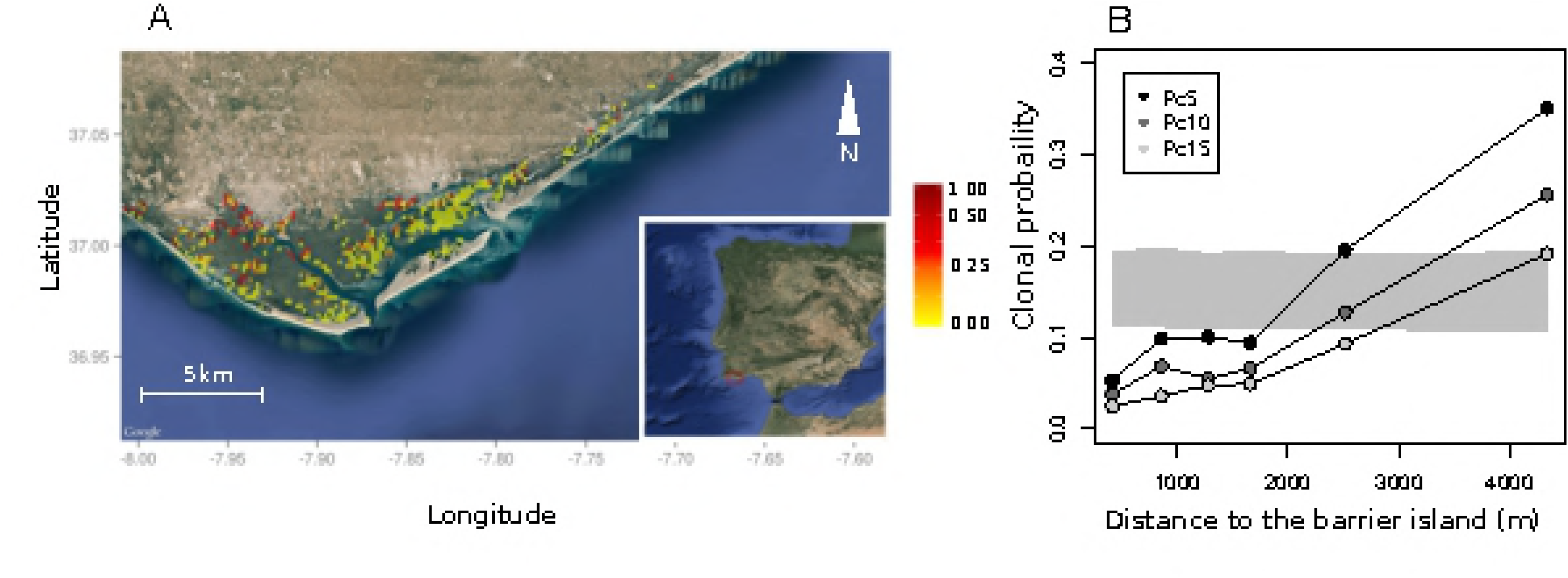
Spatial distribution of the probability of sampling a large clone of *Z. noltei*. Panel a shows the plot specific probability of sampling a clone that was observed in the sample five or more times (Pc*5*). Higher values were more frequent distant from the barrier islands. Panel b shows the spatial autocorrelograms of mean Pcn probabilities with distance from the barrier island, Pc*5*, Pc*10* and Pc*15*. The gray envelope is the 95% confidence band for the expected Pc5 under the null hypothesis of random spatial distribution, i.e., no association between sampling larger clones and distance to barrier island.

## Discussion

The spatial distribution of *Z. noltei* clones in Ria Formosa revealed multiple identical MLGs separated by tens of kilometers, despite the high genetic and genotypic diversity found. The large sample size used allowed finding for the first time several of these large clones. Given the extremely old ages found using this method, many of the largest clones sampled had estimated points of origin on land at their inferred time of clone initiation. Upon correcting our estimates for sea-level change, the estimated age of several clones within the lagoon surpassed 20, and even 30, thousand years. The estimation of replicated, extremely old organisms is on its own quite interesting and would add *Z. noltei* to the list of the oldest living organisms on the planet. However, we argue that rhizome elongation is not the most parsimonious model to explain our results. The large spatial extent covered by these seagrass clones is best explained by asexual hydrochory corroborating our previous findings on the potential for transport [46] and establishment of asexual fragments in this species [47].

The relative young age of Ria Formosa lagoon, when compared to clone ages estimated above, is evidence against a parsimonious model of clonal spread relying exclusively on rhizome elongation. The age of the Ria Formosa lagoon is still a matter of debate. Estimates vary from two thousand YBP to about six and a half thousand YBP [65,67]. Thus, any clone older than seven thousand years would not have originated inside the lagoon. Contrary to the rhizome elongation aging method, effective asexual hydrochory dispersal would hinder aging clones. However, we note that clone age can be much lower if AH dispersal proceeds in a stepping-stone manner. Support for stepping-stone AH dispersal comes from the observation of many clones with large spatial extent. If AH dispersal is the mechanism explaining the spatial extent of this many clones, then the same mechanism is likely recurrent within the clone as it expands. Although the rate of stepping stone AH dispersal is unquantifiable, our previous work with *Z. noltei*, showing long viability and high rates of successful establishment of vegetative fragments, suggests this might be more common and relatively fast than previously believed.

Mounting empirical evidence in other aquatic species, including seagrasses, supports the three necessary stages of *separation, transport* and *establishment* for successful AH. Separation of clonal modules and transport of fragments is known in many partially clonal organisms [11,14,19,21–24,68–70]. Moreover, dispersal via vegetative fragments is well documented in freshwater flora. Sand-Jensen et al. [12] found that 90% of new macrophyte patches in streams developed from the vegetative fragments that settled in a new area. In their review, De Meester et al. [16] corroborated this finding with examples from other plant and animal species. Finally, Riis & Sand-Jensen [17] followed up on this work, estimating the dispersal distance of plant fragments. In seagrasses, natural detachment, drifting and re-rooting was observed for small fragments of *Posidonia oceanica* in the Balearic Islands, Spain [40], for two other *Posidonia* species in Western Australia, with varying success [71], and for *Halophila johnsonii* and *H. decipiens* in Florida, USA [39].

As the interest in seagrass dispersal is growing in the last years [32–34,37,39,41,65,72–75] we believe that AH might be found in other species if studied at a relevant scale. AH seems to be possible for a range of seagrass species that cover a range of sizes, from some of the larger species from genus *Posidonia* [40,71] to one of the smaller species *Halophila johnsonii* [41]. Recently, an extremely large clone, spread over 47km, was found for the seagrass *Thallassia testidinum* in Florida using microsatellite analysis [9]. The authors concluded that AH dispersal was the likely process behind the propagation of this organism. Another recent study on *Zostera noltei* genetic and clonal structure, aimed at understanding the mechanism of dispersal in the Black Sea, concluded that sexual dispersal is the mechanism for long distance dispersal because no clonemates were found among populations [76]. That study provides an opportunity to compare the statistical power of individual (our study) versus population based [76] sampling designs to discover AH. Assuming that AH occurs with similar rate in Ria Formosa and Black Sea (for the sole purpose of considering statistical power) we compared the proportions of clonemates sampled at different distances in our study (Fig 2) with sample sizes and distances between populations in Jahnke et al. [76]. We estimated that the latter study would have sampled 6, 4 and 0.1 clonemate pairs at 1, 5 and 10 km distances, respectively. Although no clonemates were actually found among populations in Jahnke et al. [76], the study had small power to detect a strong AH effect, such as observed in Ria Formosa. Much smaller rates of AH would still be biological significant and undetectable under classic population based sampling designs.

Interestingly, from the other two seagrass species found in Ria Formosa, in *Cymodocea nodosa* we previously found an even larger (43km) male clone, distributed across the entire lagoon [77,78]. When we applied the aging methods used in this study to *C. nodosa*, we found that this single clone would have to be 53 000 years old (table 1). The other seagrass present in Ria Formosa, *Zostera marina*, did not show large clonal spans [79], although its patch extinction-recolonization dynamics in Ria Formosa suggests that seed dispersal via floating fragments might be an important process.

### Significance for seagrass conservation

An alarming rate of seagrass ecosystem decline has been observed in the last decades [80–84]. The rates of seagrass recovery on their own, assuming removal of the pressures that led to extinction, may be underestimated by failing to acknowledge the potential for AH dispersal. For example, in a recent synthesis on seagrass dispersal a model was used to estimate the multiple generation time that different seagrass species would take to disperse over distances ranging from meters to thousands of kilometers [6]. The time needed to disperse over 1 to 10 km distances solely by asexual reproduction (i.e., rhizome elongation) was estimated to be orders of magnitude longer (thousands of years) than by dispersing seeds (weeks). However, mounting evidence provided above supports that asexual dispersal can be as fast as sexual dispersal. We note that the only way a seagrass sexual propagule can disperse as far as an asexual one is precisely by being transported by water while coupled to a “vegetative” fragment. However, there has been far more attention to the fate of such sexual propagules than to the fate of the vegetative vector. Moreover, the potential reduction of genetic diversity in patches colonized by asexual propagules after extinction might be counteracted by the migration rate of such propagules. This could be generalized to other seagrasses if their AH dispersal is as frequent as for *Z. noltei* in Ria Formosa, like the large number of replicated spatially disjunct clones indicates. Additionally, while our focus here has been in the asexual dispersal component, floating fragments can transport maturing fertilized flowers and can release viable seeds, which can impede lost of genetic diversity. These are optimistic news for seagrass ecosystem recovery that should attract the attention of marine ecologists interested in studying the dynamics of seagrass ecosystems.

We found higher probability of sampling large clones with increasing distance from barrier islands, assumed to be a source of physical disturbance through increased burial [66]. This association can be the result of two non-mutually exclusive processes linked to the disturbance regime - different clone survival and different allocations to sexual and asexual reproductive components. Survival and longevity might be affected by different habitat stability, with increased stability in the inner area of the lagoon allowing established clones to survive longer and thus grow larger. In contrast, in the disturbed areas meadows are frequently buried by increased sedimentation associated to the barrier island inlets and their migration. Survival of clones in such a habitat is likely lower, as the conditions are less favorable due to higher burial rates, increased turbidity and disturbance frequency.

Simultaneously, higher allocation to sexual propagation is shown for seagrasses in disturbed habitats. Gallegos et al. [85] showed four-fold increase in flowering for *Thalassia testudinum* after the disturbance cause by a hurricane passage. Looking at *C. nodosa* response to sediment dynamics via sand dune migration, Marbà & Duarte [86] noted overall higher flowering frequency in the studied disturbed meadow, in comparison to the data for non-disturbed meadows. In particular, fragments which were buried just before the flowering season showed highest frequencies of flowering. In 72% of studies on this topic, seagrasses responded to disturbance by increasing reproductive effort by 4-fold [87]. The same was shown for terrestrial grasses over 30 years ago [88]. Thus, our study indicates some interesting directions for future experimental work, tackling the relative importance of variable clone survival, as suggested by Bricker et al. [9], and variable allocation to sexual and asexual reproduction in association with the disturbance regime.

## Acknowledgements

Authors wish to acknowledge S. Almeida, D. Paulo and other colleagues for field assistance and F. Barbosa for carefully reviewing the manuscript.

## Supporting information

S1 File. Estimating clonal age using rhizome elongation rates and paleoceanographical sea-level change

**S1 Fig. Seagrass age estimation integrating paleoceanographical sea-level change**. Example of one iteration of *Zostera noltei* age estimation based on horizontal rhizome elongation rate and paleoceanographical sea-level reconstitution of the Ria Formosa lagoon area. A and B are locations of ramets belonging to the same clone. O_0_ is the initial point of origin for the clone, placed midway between points A and B. Assuming rhizome elongation only, the time to necessary for the clone to grow from O_0_ to A and B is 17 710 years. At that period, 17 000 years before present, the sea level was 120 m lower than in the present (indicated by by the red dotted line). Thus, the point of origin is corrected to O_1_, the coast line at that period, at an intermediate distance from A and B. The clone age t_1_ is updated with the time necessary to grow from the corrected O_1_ to sample sites A and B, here 32 352 years. Other discontinuous lines represent the coastline location during different periods in the past.

S2 File. Disturbance and clone size associations

